# Deciphering interaction syntax via decoupling intrinsic lineages and niche pressure

**DOI:** 10.64898/2026.05.18.725889

**Authors:** Qirui Guo, Wenhao Zhong, Zhiwei Zeng, Qing Nie, Peijie Zhou, Lei Zhang

**Author notes:** These authors contributed equally to this work (Qirui Guo, Wenhao Zhong, Zhiwei Zeng).

## Abstract

Spatial transcriptomics enables the mapping of gene expression within intact tissues, yet a fundamental gap remains between knowing where cells are and understanding how they interact. A cell’s measured transcriptome reflects both its intrinsic lineage identity and niche pressure. Here we introduce TRINUS, a self-supervised model that deciphers interaction syntax by generative decoupling of a cell’s intrinsic lineage identity from the extrinsic niche pressure. TRINUS maintains a library of context-free cell prototypes to isolate lineage identity while modeling cooperative interaction dependencies among neighbors. We validated TRINUS on synthetic datasets with known interaction logic and benchmarked it against existing methods with superior performance in cell clustering and spatial domain detection. Applied across diverse platforms and biological systems, TRINUS resolves multi-level interaction syntax and maps tissue-wide interaction patterns in colorectal cancer, and identifies stage-specific signaling dependencies and time-dependent receptor windows during mouse organogenesis. We also show TRINUS’s bidirectional *in silico* engineering capability in the ovarian tumor microenvironment, where forward perturbation revealed subtype-specific macrophage immunosuppressive programs via virtual transplantation and inverse design identified molecular modifications in macrophages predicted to rescue adjacent T-cell function. Collectively, TRINUS provides a practical tool for interaction syntax discovery and predictive tissue engineering on spatial transcriptomics data.

## Introduction

From the orchestration of embryogenesis to the dysregulation of the tumor microenvironment, tissue function is an emergent property arising not just from the presence of specific cell types, but also from the complex network of communication that connects them. The advent of single-cell Spatial Transcriptomics (ST)^1–5^ has enabled mapping of gene expression profiles to spatial coordinates, yet a fundamental gap remains between knowing where cells are and understanding how they interact. Bridging this gap requires identifying the underlying interaction syntax by which neighboring cells selectively modulate each other’s phenotypes. Extracting these rules is challenging because interactions are combinatorial, context-dependent, and inseparable from each cell’s intrinsic identity in raw measurements, demanding computational approaches that move beyond spatial co-expression analysis.

Several computational strategies have been developed toward this goal. Ligand-receptor communication inference methods^6–12^ (e.g., *COMMOT*^8^, *NicheNet*^6^) quantify intercellular signaling by scoring the co-expression of known molecular pairs. These methods have been valuable for mapping communication landscapes, but their reliance on biological priors constrains them to characterized interactions in an unsupervised manner, potentially limiting the discovery of novel or context-dependent interaction rules.

Another data-driven strategy uses graph neural networks (GNN)^13–21^ (e.g., *STAGATE*^13^, *SpaGCN*^15^) to learn cell representations that incorporate spatial context. By aggregating information from neighboring cells, these models effectively denoise spatial measurements and refine cell-level representations for downstream clustering. However, these methods fuse the cell and its context into a single embedding. This makes it difficult to distinguish expression changes driven by the microenvironment from those reflecting intrinsic lineage, which is central to isolating interaction syntax.

To make this distinction, a recent class of methods explicitly decomposes cell states into intrinsic and extrinsic components. Residual-based methods^22–25^ such as *GITIII*^24^ estimate a cell’s intrinsic baseline from the mean expression of its annotated cell type and attribute the remainder to environmental influence. While intuitive, this strategy depends on pre-defined cell-type labels and assumes that population averages adequately represent intrinsic identity, which can introduce bias, particularly for heterogeneous or transitional cell states. Latent factorization methods^26–33^ (e.g., *NicheCompass*^28^, *SIMVI*^26^, *Decipher*^27^) avoid this dependency by learning separate latent spaces through parallel encoders. However, these methods generally treat the neighborhood as an unordered collection of cells, considering each neighbor’s contribution independently. This makes it challenging to capture the cooperative and combinatorial nature of cellular interactions, for instance, cases where a signaling outcome depends not only on which cell types are present but on whether those cells are actually interacting.

Beyond descriptive analysis, an important direction is to move from observation toward intervention, not only predicting how tissues respond to perturbations but also designing microenvironmental changes that produce desired cellular outcomes. While existing methods have enabled forward perturbation to simulate the resident tissue’s response to specific modifications on observational spatial transcriptomics data (e.g. *MintFlow*^33^, *Celcomen*^29^) or on spatially resolved perturbation screens (*CONCERT*^34^), the complementary inverse problem, i.e. identifying the minimal spatial remodeling required to drive a specific phenotypic transition, remains largely unaddressed^35,36^. Solving this inverse problem could substantially reduce the experimental search space for tissue engineering, transforming a combinatorially vast space of possible interventions into a tractable set of prioritized hypotheses for validation.

To address these limitations, we introduce TRINUS (**T**riadic **R**epresentation of **I**nteracting **N**iches for **U**nderlying **S**yntax), a self-supervised model that deciphers interaction syntax through the generative decoupling of a cell’s intrinsic identity from extrinsic niche pressure. TRINUS incorporates three key design principles. First, to capture the cooperative and combinatorial nature of cellular interactions, we move beyond standard graph aggregation to explicitly model how the relationship between two cells depends on their mutual neighbors (triangle multiplicative attention^37,38^). This enables the model to distinguish active interaction syntax from coincidental spatial co-occurrence. Second, to isolate intrinsic identity without relying on prior cell type annotations, we build a dynamically updated library of context-free cell states (Vector-Quantized (VQ) codebook^39–41^). Each cell’s observed expression is decomposed into a discrete lineage component matched to this library and a continuous interaction-induced deviation, enabling label-free quantification of niche influence. Finally, the joint representation of cellular states and interaction patterns establishes a differentiable space that supports bidirectional *in silico* engineering, allowing both the simulation of tissue responses to virtual perturbations and the identification of minimal neighborhood modifications required to drive a target phenotypic transition.

We first validated TRINUS on synthetic datasets with known interaction rules, confirming its ability to recover non-linear and cooperative logic that standard graph neural networks are challenging to capture. Through comprehensive benchmarking on real-world spatial transcriptomics data, TRINUS outperformed existing methods in both cell clustering and spatial domain detection, accurately resolving fine-grained cellular and niche heterogeneity. We then applied TRINUS across diverse platforms and biological systems with demonstrated utility in decoding context-specific signaling patterns and guiding in silico perturbations. In colorectal tumor tissue (Visium HD), the model resolved multi-level heterogeneity and mapped tissue-wide interaction patterns through an interaction compass, revealing recurring communication motifs and their associated signaling efficacy. In mouse embryonic development (Stereo-seq, E9.5– E16.5), TRINUS identified stage-specific signaling dependencies, distinguishing transient developmental cues from constitutive maintenance signals. In the ovarian tumor microenvironment (Xenium 5K), we demonstrated bidirectional *in silico* engineering by performing virtual macrophage transplantations and inverse design experiments to identify molecular modifications predicted to rescue T-cell function. We further show that TRINUS can enhance single-cell foundation models by providing spatial interaction context. Collectively, these results demonstrate that TRINUS provides a practical and generalizable tool for interaction syntax discovery and predictive tissue engineering.

## Results

### TRINUS decodes interaction syntax through generative decoupling

To investigate the interaction syntax underlying the spatial transcriptomics data, we developed TRINUS, a deep generative model that treats tissue function as a system of interactive components between both cell lineages and their interactions. Given spatial transcriptomics data, TRINUS models tissue as a system of proximal collections of interacting cells called niches. From the input niche, TRINUS initializes cell embeddings that encode cellular transcriptomic states and interaction embeddings that capture the spatial proximity and expression correlation between cells (Fig. 1a(i)). These streams define the input for our core architecture, the stacked NicheLayer (Fig. 1a(ii)). Unlike standard GNNs that aggregate neighborhoods into isotropic scalars, this module iteratively refines both embeddings using triangular multiplicative updates and axial attention^37,38^. The dual-stream architecture allows the model to capture higher-order interaction patterns that are invisible to standard node/edge information aggregation, transforming the initial inputs to informative context-aware cell embedding and interaction embeddings.

**Fig 1.**
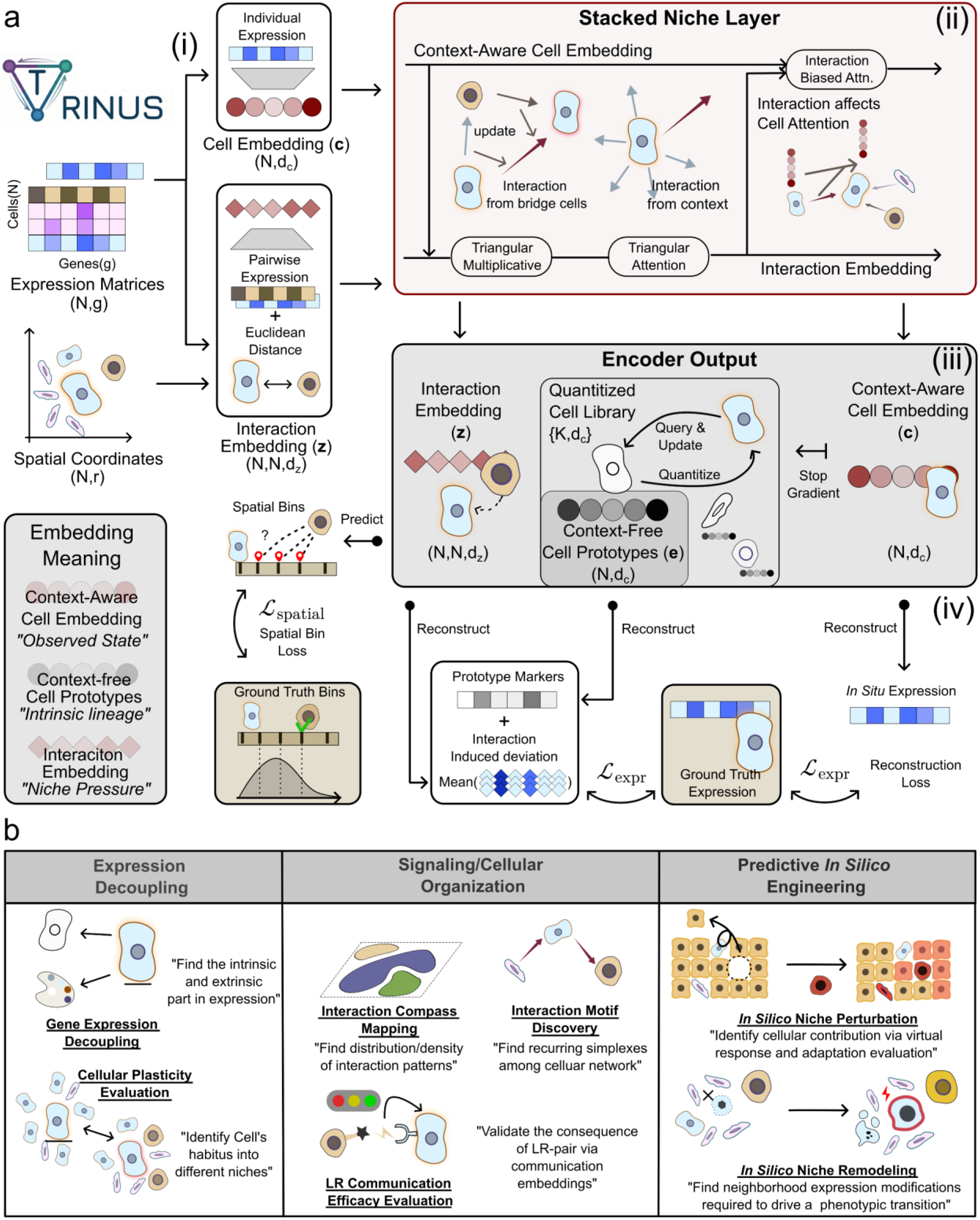
The TRINUS framework for deciphering interaction syntax. (a) Model Architecture. (i) Initialization of context aware cell embeddings (c) and interaction embeddings (**z**) from spatial transcriptomics input niches. (ii) Iterative refinement of both embeddings via stacked NicheLayers to facilitate information exchange between cellular and interaction representations. (iii) Mapping of continuous cell representations to discrete, context free prototypes (**e**) via a Vector Quantized (VQ) bottleneck. (iv) Self-supervised decoupling objectives, comprising a hurdle loss for cell reconstruction, a niche residual loss predicting interaction-induced expression deviations from prototype markers, and a spatial binning loss for interaction distance awareness. (b) Downstream applications enabled by generative decoupling. (Left) Expression decoupling into intrinsic and interaction-induced components for the evaluation of in situ cellular plasticity. (Middle) Characterization of cellular interaction organization, including the discovery of high-order communication motifs, evaluation of communication efficacy, and spatial mapping via an interaction compass. (Right) Bidirectional in silico tissue engineering, enabling forward perturbation via virtual transplantation and inverse niche remodeling to drive target phenotypes.

To isolate intrinsic identity, TRINUS builds up a vector quantized cell library that maps the continuous context-aware representations onto a discrete library of context-free prototypes. Such a process filters out the continuous fluctuations introduced by the microenvironment, retaining only the canonical lineage identities. Finally, the model simultaneously builds three embeddings (Fig. 1a(iii)): context aware cell embeddings, context-free prototypes and interaction embeddings. To further decipher cellular states and syntax of interaction, we constructed comprehensive training objectives for TRINUS (Fig. 1a(iv); **Methods**). By leveraging interaction embedding to reconstruct interaction-induced deviation, residual between observed gene expression and decoded intrinsic markers from prototypes, TRINUS explicitly captures the effects that the microenvironment exerts on cellular identity.

The embeddings in TRINUS enable three categories of downstream applications (Fig. 1b; **Methods**). At the cellular level, TRINUS performs expression decoupling and unsupervised cell plasticity evaluation, allowing users to attribute specific gene expression programs to either lineage fidelity or niche adaptation without requiring ground-truth labels. At the tissue level, leveraging the dense interaction embeddings, TRINUS deciphers signaling and cellular network organization among tissue. This ranges from quantifying the efficacy of ligand-receptor communication, discovering recurring high-order interaction motifs and mapping the vector fields of tissue-wide communication pattern via an interaction compass. Finally, TRINUS unlocks both forward and inverse *in silico* engineering. Functioning as a differentiable simulator, TRINUS not only allows forward perturbation to predict the niche response to virtual transplantation and adaptation of transplanted cells to new niche. This predictive capability also enables inverse remodeling, where the model seeks to identify the minimal neighborhood expression remodeling required to drive a desired phenotypical transition.

### Validation and benchmark of TRINUS’s generative decoupling function

To ensure that TRINUS accurately learns the governing rules of interaction rather than memorizing statistical correlations, we first verified our key architectural hypotheses on controlled synthetic datasets and benchmarking on the real-world dataset. We first illustrate the challenges of standard GNNs using two synthetic logic gates (Supplementary Fig. 1). In the first scenario where a receptor is activated only by spatially distinct senders but blocked if senders contact with each other (lateral interference, Supplementary Fig. 1a), standard GNNs struggle to identify such an interaction pattern due to their additive star topology aggregation. In contrast, TRINUS utilizes triangular attention to model lateral inhibitory edges, functioning as a non-linear gate to correctly predict the blocked state. Similarly, in the second scenario where a receptor is blocked only when surrounded by a dense community of senders (enclosure suppression, Supplementary Fig. 1b), our model distinguished active cliques from dispersed neighbors by learning higher-order interaction dependencies.

We further validated the expression decoupling performance (Supplementary Fig. 2) against reference-dependent residual-based methods (e.g., GITIII^24^). On synthetic data governed by known rules, GITIII struggled to construct precise intrinsic identity profiles via static population means, which consequently reduced the accuracy of its niche pressure estimations. Conversely, TRINUS accurately recovered both lineage markers and niche pressures. This capability of learning interaction syntax was further investigated via synthetic *in silico* perturbation tests with pre-defined interaction rules (Supplementary Fig. 3). In forward perturbation tests (Supplementary Fig. 3a), TRINUS correctly predicted the specific gene expression shifts in response to neighbor replacement. Meanwhile, in the inverse design task (Supplementary Fig. 3b), the model found the desired programmed spatial dependencies, identifying the exact expression remodeling responsible for triggering a specific phenotype transition. This confirms that TRINUS does not merely hallucinate solutions but accurately retrieves the underlying interaction logic governed by the ground truth.

Having mechanically verified these architectural designs, we further investigate whether the dual stream architecture would benefit context-aware cell embedding on a human Xenium 5k COAD dataset (Fig. 2a). This was confirmed empirically: when benchmarked against Banksy^42^ and STAGATE^13^, TRINUS achieved superior performance across NMI, ARI, homogeneity, and completeness metrics (Fig. 2c, **Supplementary Notes**; **Methods**), with distinct improvements in biologically heterogeneous regions (Fig. 2b). Notably, the model leveraged its decoupling capacity to resolve fine-grained heterogeneity, stratifying epithelial cells that were originally annotated as a single broad class into five distinct subclusters representing metastable states (Fig. 2d-g), including specific stem-like and metabolic phenotypes confirmed by differential expression.

**Fig 2.**
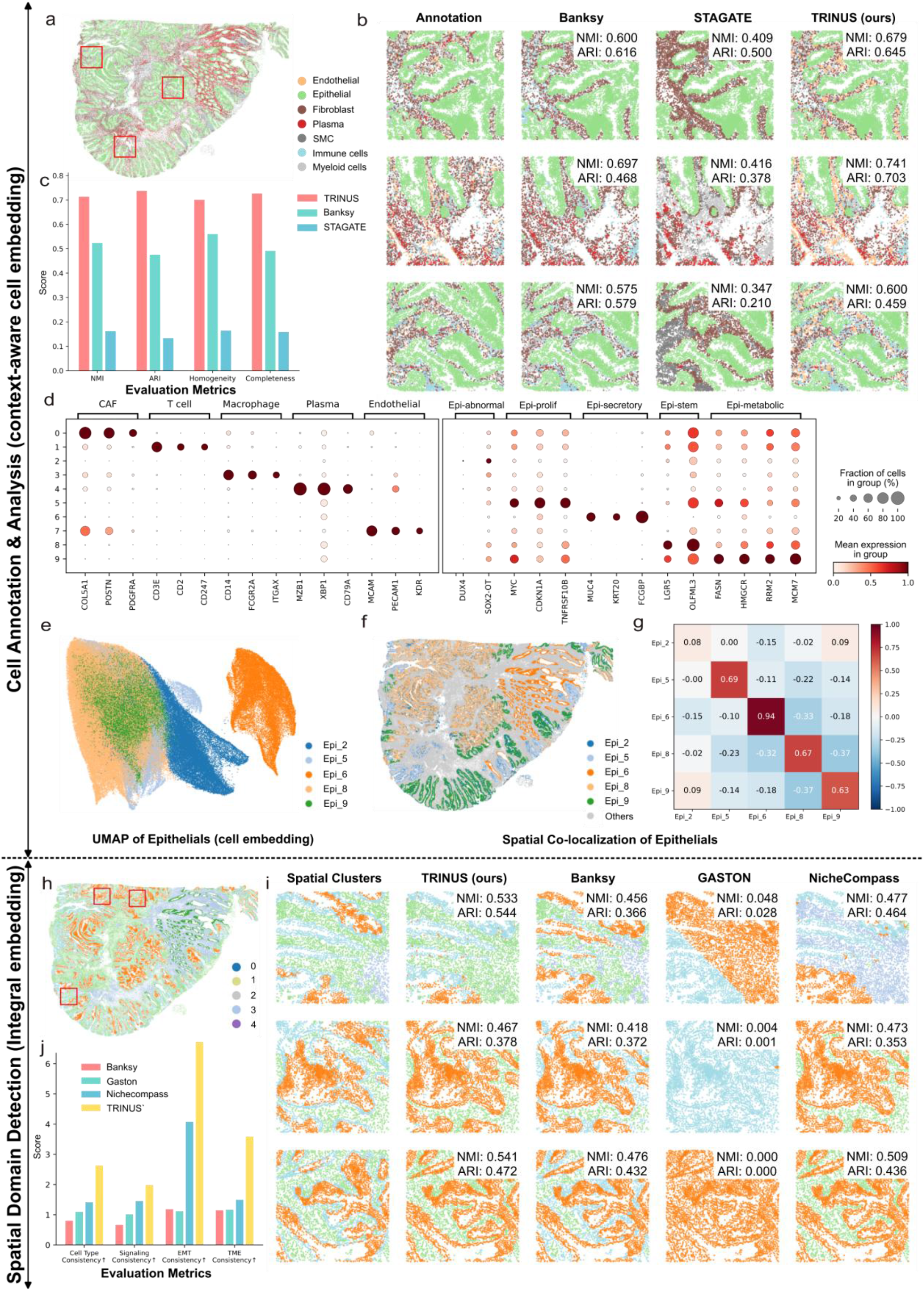
Benchmarking TRINUS on cell clustering and spatial domain detection. (a) Spatial overview of the human colon adenocarcinoma (COAD) Xenium 5K dataset, annotated by cell type. (b) Comparison of cell clustering spatial maps generated by Banksy, STAGATE, and TRINUS against ground truth within the regions highlighted by red rectangles in (a). (c) Evaluation metrics (NMI, ARI, Homogeneity, and Completeness) for the clustering of non-epithelial cells across the benchmarked methods. (d) Dot plot of marker gene expression across TRINUS-identified major cell types (CAF, T cell, macrophage, plasma, endothelial) and high-resolution epithelial subtypes (abnormal, proliferative, secretory, stem-like, metabolic). (e) UMAP visualization illustrating the distinct epithelial states identified by TRINUS. (f) Spatial distribution of different epithelial subtypes across the tissue. (g) Spatial self-regression matrix of the epithelial subtypes. (h) Spatial overview of the COAD dataset annotated by structural spatial domains. (i) Comparison of spatial domain detection results from Banksy, GASTON, NicheCompass, and TRINUS within the regions highlighted by red rectangles in (h). (j) Overall evaluation metrics for spatial domain detection across the benchmarked methods

Finally, we evaluated the model’s capacity to reconstruct tissue architecture by integrating cell and interaction embeddings for spatial domain detection. Compared with Banksy^42^, GASTON^43^, and NicheCompass^28^ (Fig. 2h), TRINUS demonstrated a superior ability to resolve histologically complex regions (Fig. 2i), yielding clusters with high biological coherence (Fig. 2j). The identified domains correspond to functionally specialized compartments (Supplementary Fig. 6), including: an immune-excluded tumor core (cluster 3) dominated by stem-like epithelial cells; a metabolic transition zone (cluster 1); a stromal immune-rich region (cluster 0); a secretory immune interface (cluster 4) featuring triadic interactions between secretory cells and cDC2+ APCs; and a tertiary lymphoid structure (cluster 2). The precise segmentation results of TRINUS capture the underlying tissue structure beyond simple proximity metrics. Sensitivity analysis revealed that while cell annotation metrics peaked with higher weights on cell embeddings, domain detection required balanced contributions from interaction embeddings (Supplementary Fig. 4). This trade-off suggests that a comprehensive definition of functional tissue niches benefits from integrating cellular identity with the structural context of the microenvironment. Overall, these results suggest that TRINUS could identify interaction syntax among complex tissue structures by capturing the dependencies between cellular identity and niche pressure.

#### TRINUS decouples lineage identity and niche pressure in colorectal cancer

To investigate the capacity of TRINUS to resolve the underlying syntax governing cellular communities, we applied the model to high-resolution spatial transcriptomics (VisiumHD) profiles of colorectal tumor tissue (Fig. 3a). By training the model to reconstruct observed gene expression as a composite of discrete intrinsic markers and continuous interaction-induced deviations, TRINUS learned a library of context-free cell prototypes (Fig. 3b, Supplementary Fig. 7). These prototypes serve as anchors in embedding space and enable the prediction of intrinsic markers with high lineage fidelity (Fig. 3c, Supplementary Fig. 8-10). The model resolved a complex cellular hierarchy, distinguishing inflammatory fibroblasts (P33; CXCL12, SFRP1) from pathological, desmoplastic cancer-associated fibroblasts (P23; CTHRC1, PDPN), consistent with the established iCAF–myCAF differences^44^. It also simultaneously resolved the tumor heterogeneity into functionally distinct identities: a differentiated tumor population retaining colonocyte identity (P36; SLC26A3^45^), invasive states (P8; MAP3K20^46^), and a stem-like niche (P39, LEFTY1^47^), alongside an immune compartment (P48; CXCL13,MS4A1^48^). These assignments were corroborated by differential expression analysis (Supplementary Fig. 11), indicating that the prototype learning strategy effectively segregates lineage-specific gene expression.

**Fig 3.**
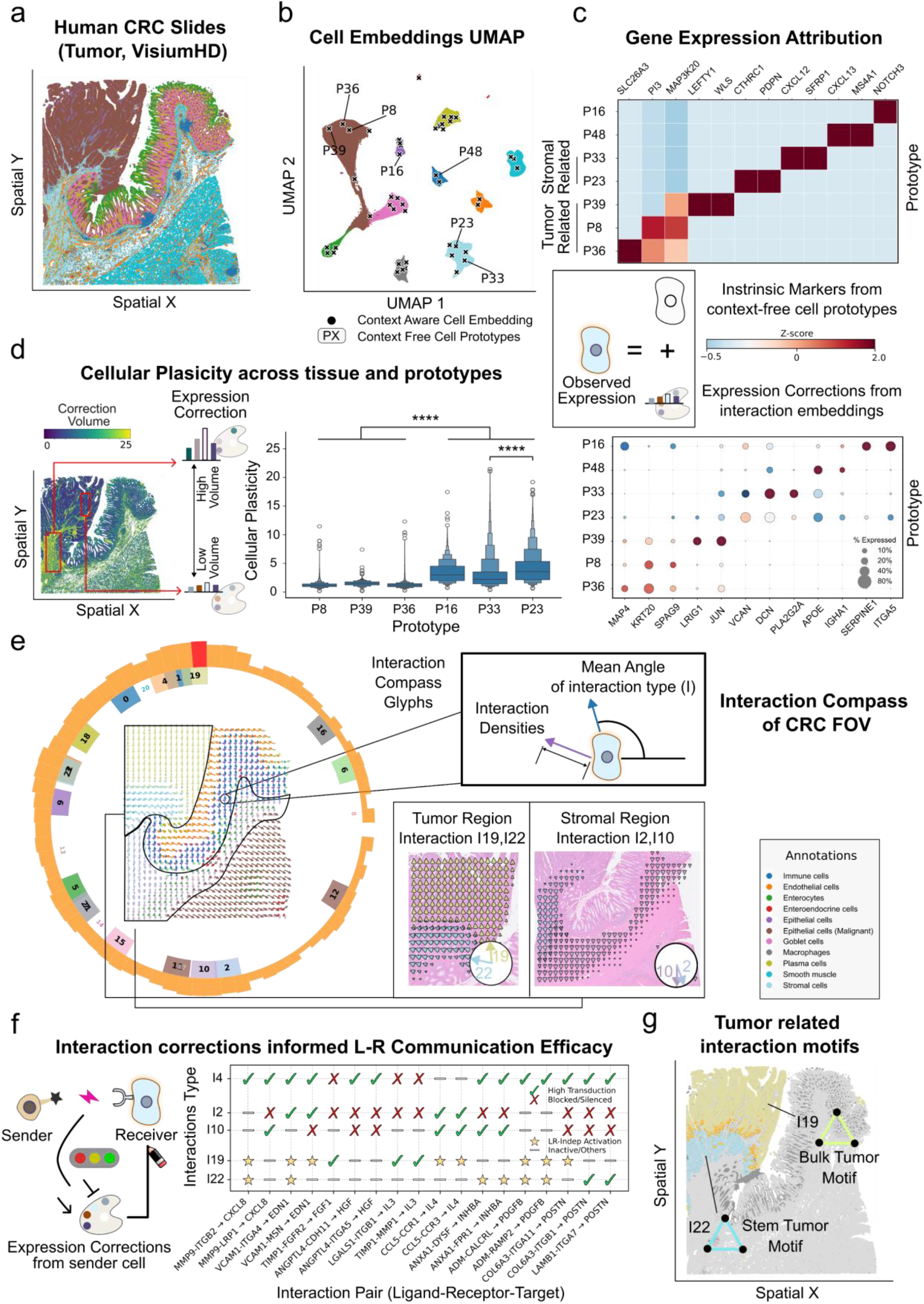
TRINUS resolves multi-scale interaction syntax and cellular plasticity in colorectal cancer. (a) Spatial transcriptomics profiling of human colorectal cancer (CRC) tissue using the VisiumHD platform. (b) UMAP of context-aware cell embeddings (colored dot) and context free cell embeddings (PX). (c) Decoupling of observed gene expression into intrinsic marker profiles derived from context-free prototypes (top) and interaction-induced expression corrections across distinct prototypes (bottom). (d) Spatial mapping (left) and prototype-specific distributions (right) of cellular plasticity, defined as the overall magnitude of interaction-induced expression deviation. (e) Visualization of the interaction compass derived from TRINUS interaction embeddings. Glyphs encode specific interaction types via azimuth and local signaling density via magnitude. Callouts show the macroscopic angular distribution of interaction vectors and the microscopic vector fields consisting of specific tumor (I19, I22) and stromal (I2, I10) related interaction types. (f) Evaluation of ligand-receptor (LR) communication efficacy. (Left) Illustration of the integration of structural LR co-expression potential with model-inferred, interaction-induced deviations. (Right) Communication efficacy profiles across distinct interaction types, classifying specific signaling axes as actively transduced, blocked, or LR-independent. (g) Spatial distribution of identified tumor-related interaction motifs (I19 Bulk tumor motif and I22 Stem tumor motif).

Beyond lineage identity, TRINUS explicitly models the niche pressure via the interaction embedding. This dense representation allows the model to infer the specific interaction-induced deviation required to drive a cell’s intrinsic state to its observed phenotype. We defined cellular plasticity as the magnitude of this inferred deviation vector (**Methods**). Mapping this metric revealed the landscape of differential adaptability across the tissue. Supporting lineages, specifically endothelial cells (P16) and fibroblasts (P23, P33), exhibited higher plasticity scores compared to the relatively stable tumor core (P8, P36) (Fig. 3d). Furthermore, the specific gene content of these deviations aligned with context-specific functions rather than random noise (Fig. 3c, Supplementary Fig. 12). For instance, while P16 endothelial deviations were enriched for remodeling factors SERPINE3 and ITGA5^49^, P23 fibroblasts showed specific deviation towards the malignancy-supporting VCAN^50^ state. The alignment of these expression deviations with known remodeling factors suggests that the interaction embedding captures context-specific shifts, providing a quantitative metric for distinguishing intrinsic stability from niche-associated adaptation.

To further investigate the syntax of tissue-wide interaction, we mapped the spatial distribution of the learned interaction embeddings via an interaction compass (Fig. 3e, Supplementary Fig. 13; **Methods**). In this vector-field representation, we introduced glyphs whose azimuth encodes the specific interaction type (*I*) and the magnitude represents the local signaling density. Interestingly, the interaction compass revealed a topological divergence unobservable in gene expression space alone: stromal regions displayed a coherent flow of spatially adjacent vectors (dominated by Types I2 and I10), whereas the tumor region was characterized by the orthogonal intersection of distinct vector fields (Types I19 and I22). These coherent vector fields suggest that the interaction compass reveals tissue organization features that are not apparent from cell type distribution alone, mapping the spatial continuity of interaction logic.

We next leveraged the interaction embeddings to bridge the gap between structural connectivity and functional outcome. While standard cell-cell communication analysis relies on static Ligand-Receptor co-expression, TRINUS integrates this structural potential with the model-inferred response (interaction-induced deviations) to evaluate communication efficacy (Fig. 3f, Supplementary Fig. 14; **Methods**). This analysis highlights the functional consequence of signaling by distinguishing where signals are actively transduced versus where they are silenced. Evaluating these efficacy profiles suggests distinct regulatory logic across compartments. For instance, stromal interactions displayed a mix of high transduction and blocked states, whereas tumor-associated interactions (I19, I22) were dominated by LR-independent activation. These distinct responses suggest the underlying interaction modes, identifying that inflammatory networks (I10) exhibited high transduction efficacy for MMP9-to-CXCL8^51^ signaling, whereas pathological networks (I2) showed suppressed efficacy for this axis, coinciding with VCAN-driven desmoplasia. Conversely, the tumor’s autonomous interactions (I19) drove a survival TIMP1^52^ signaling axis, which was distinct from the mechanosensing zones (I22) anchored to integrin and POSTN^53^ signaling. By distinguishing potential from response, the interaction embedding enables the isolation of active regulatory modes from background expression.

Having established the molecular syntax of individual interaction types, we sought to reconstruct the multi-cellular architecture formed by their assembly. We analyzed the composition of high-order communication motifs (Fig. 3g, Supplementary Fig. 15; **Methods**) to determine whether the interaction types correspond to spatial neighborhoods. By extracting recurring simplexes from the interaction graph, we found that the autonomous interaction motifs (I19-motifs) were structurally composed of P36/P8 bulk tumor cells, while the mechanosensing motifs (I22-motifs) specifically anchored P39 stem-like cells to the matrix. Similarly, in the stroma, motifs segregated by fibroblast subtype, linking the laminar flow of the interaction compass directly to spatially coherent cell neighborhoods. The correspondence between these interaction simplexes and distinct histological zones indicates that triangular attention captures the high-order geometric dependencies defining local tissue architecture.

In summary, the recovery of distinct cellular prototypes and the mapping of spatial interaction patterns suggest that tissue phenotypes can be resolved into intrinsic and extrinsic components. By decoupling lineage Identity and niche pressure for resolving tissue heterogeneity, TRINUS enables a multi-scale characterization of the tumor microenvironment, linking individual plasticity profiles to collective signaling topologies.

#### TRINUS enables bidirectional *in silico* engineering of macrophages in ovarian tumor microenvironments

Having shown that TRINUS decouples intrinsic cellular identity from niche pressure, we leveraged its generative capability to perform *in silico* tissue engineering. We focused on the macrophage-mediated regulation of T-cell exclusion in the ovarian tumor microenvironment. While Tumor-Associated Macrophages (TAMs) are known drivers of T-cell exclusion, the interaction syntax among immune-dysfunction niches remains difficult to decode.

To investigate such immune barrier, we began with forward perturbation to show the regulatory consequences of the macrophage’s infiltration. We performed virtual transplantation experiments, digitally excising stress-induced fibroblasts and replacing them with either M1-like (inflammatory; SLC2A1,IFIT1,HK2) or M2-like (exclusionary; VSIG4,CD163,MRC1) macrophages^54–56^ identified by the two distinct context free prototypes, and focused on altered embedding or reconstructed niche expression from TRINUS (Fig. 4a,b, Supplementary Fig. 16,17; **Methods**). We also constructed control niches that replace the fibroblasts with other stress-induced fibroblasts for a niche stability baseline. This perturbation allowed us to probe the reciprocal syntax of interaction: specifically, how the resident niche responds to the transplant, and how the transplant adapts to the new niche.

**Fig 4.**
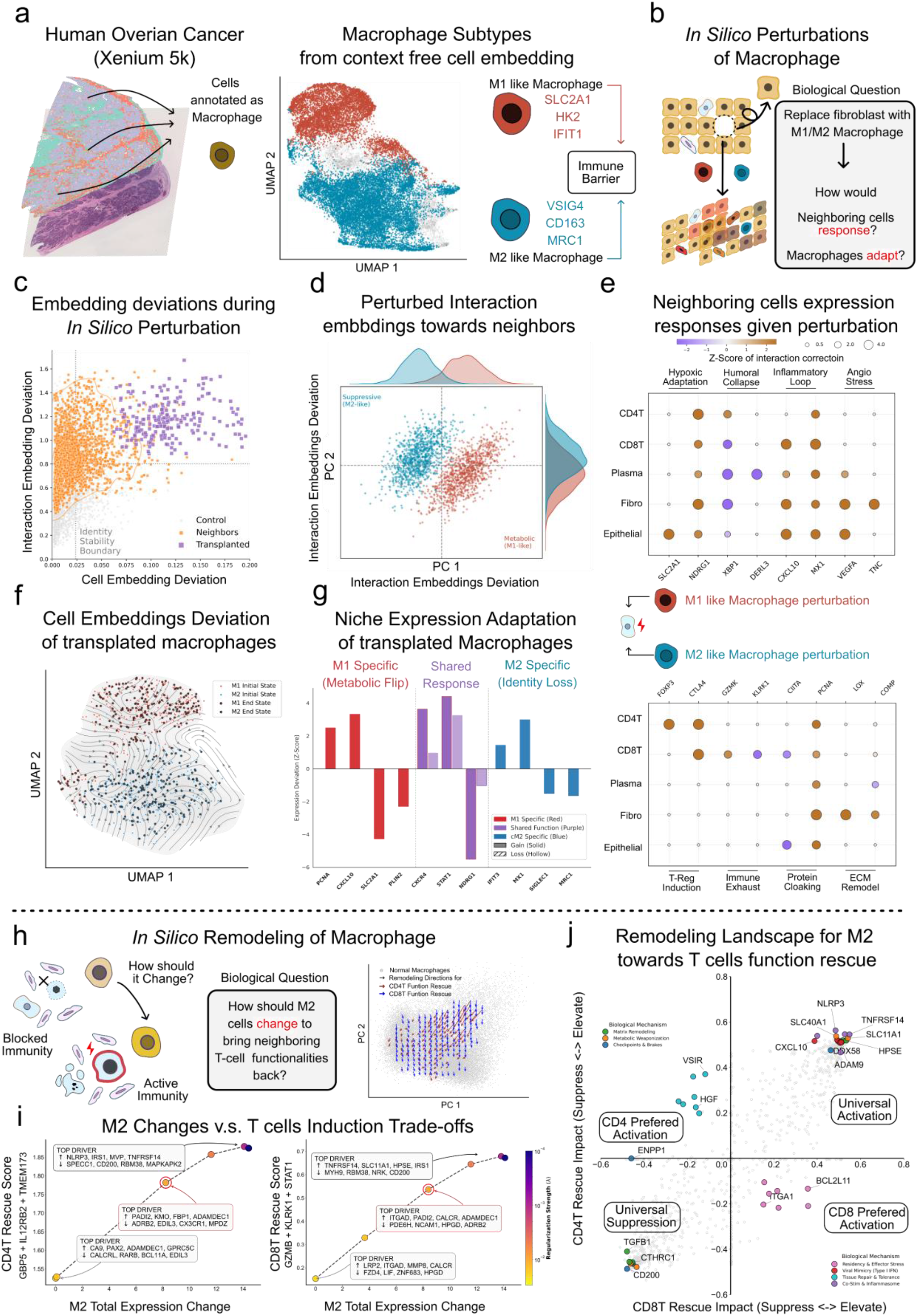
TRINUS enables bidirectional in silico engineering to decode macrophage-mediated immune barriers in ovarian cancer. (a) (left) Spatial transcriptomics profiling of the ovarian tumor microenvironment using the Xenium 5K platform. (right) UMAP of TRINUS context-aware cell embeddings for annotated macrophages, colored by assignment of context-free cell prototypes. (M1-like, red; M2-like, blue) (b) Illustration of the in silico perturbation experiment in which stress-induced fibroblasts were replaced by M1-like or M2-like macrophages to model niche responses and cell adaptation. (c) Magnitude of structural perturbation in the TRINUS latent space. The plot maps the norm of cell embeddings delta for each cell in the region after perturbation (x-axis) against the norm of the maximum interaction embedding changes among all present interactions in each cell’s niche (y-axis). (d) PCA visualization of interaction embedding deviations between transplanted macrophages and their neighbors. (e) Niche-wide gene expression response to M1-like (top) and M2-like (bottom) virtual transplants. Dot size and color indicate the relative magnitude and direction of expression changes (control-calibrated two-sample z-scores) across diverse neighboring cell types. (f) Stream-plot of context-aware cell embeddings change of transplanted macrophage. (g) Reconstructed expression adaptation of transplanted macrophages. (h) Illustration of in silico niche remodeling experiments of M2 macrophages to rescue T cell functions. (left) This niche remodeling aims to identify the minimal transcriptional changes in suppressive M2 macrophages required to maximize functional markers in neighboring CD8+ and CD4+ T cells. (right) Remodeling trajectories of M2 macrophages in PCA space. (i) Identified remodeling genes in M2 macrophages under limited move constraints. (j)Specific gene expression remodeling required in the M2 macrophage to simultaneously rescue CD8+ T cells (x-axis; left: suppress, right: elevate) and CD4+ T cells (y-axis; bottom: suppress, top: elevate).

We first analyzed the structural impacts of this perturbation in the embedding space. The model produced a hierarchical distribution of embedding deviations given perturbed niches (Fig. 4c). Resident cells in control experiments (fibroblast-fibroblast replacement, gray) exhibited negligible embedding shifts, showing model stability. In contrast, neighbors of the macrophage transplants (orange) showed significant deviations specifically in their interaction embeddings, indicating a rewiring of interactions. Notably, the transplanted macrophages themselves (purple) exhibited the highest magnitude of deviation, suggesting they were undergoing substantial adaptation to their new environment. PCA analysis of these embedding deviations revealed a clear bifurcation between M1- and M2-induced niches (Fig. 4d). The separation of these clusters implies that the interaction embeddings capture phenotype-specific motifs distinct from generic perturbation artifacts.

The biological consequence of the perturbation was decoded into specific gene expression signatures (Fig. 4e, Supplementary Fig. 18-20). As shown, we found that M1-like transplants induced changes across multiple cellular programs: aside from inducing an interferon alarm (MX1,CXCL10^57^), they also activated a metabolic shift in which tumor cells upregulated SLC2A1 amidst universal hypoxic stress (NDRG1^58^), accompanied by downregulation of UPR master regulators in plasma cells (XBP1, DERL3^59^), suggesting compromised secretory capacity. Conversely, M2-like transplants engineered a dual-layer exclusion barrier. These transplants shifted CD4+ T-cells toward regulatory states (FOXP3+^60^,CTLA4+^61^) and CD8+ T cells toward a low-cytotoxicity, pro-inflammatory effector memory state (CTLA4+^61^,KLRK1-^62^,GZMK+^63,64^). They also led to molecular cloaking by suppressing CIITA^65^ in epithelial cells while simultaneously fortifying the physical barrier via fibroblast-mediated fibrosis (LOX^66^). These niche responses characterized M1 and M2 macrophages as distinct organizing centers that exhibited divergent metabolic and physical regulatory modes within the perturbed microenvironment.

We further investigated the adaptation of transplant to show the model’s grasp of reciprocal constraints. By visualizing the change of the context-aware embeddings (Fig. 4f), we observed that macrophages drifted along distinct trajectories. M1 adaptation was characterized by the specific loss of the lipid-loaded, glycolytic phenotype (PLIN2, SLC2A1^54^), together with the acquisition of proliferative capacity (PCNA^67^; Fig. 4g, Supplementary Fig. 20). Similarly, M2 macrophages stripped of their immunosuppressive niche lost their resident identity markers (SIGLEC1, MRC1^55,56^) and rapidly sensed the inflammatory tone (IFIT3, MX1^57^). Notably, shared downregulation of NDRG1^59^ confirms that the model correctly identifies niche-imposed hypoxic constraints. These adaptations suggest that while the transplants actively remodeled the niche, they were simultaneously constrained by environmental pressure.

Having investigated these regulatory rules via forward perturbation, we then solved the inverse remodeling problem, i.e., identifying the minimal molecular intervention upon suppressive M2 macrophages required to drive neighboring T-cell functional recovery (Fig. 4h). We defined a rescue objective maximizing cytotoxic markers (GZMB^64^, KLRK1^62^,STAT1^68^) for CD8+ T cells and Th1 recruiting markers (IL12RB2, GBP5^69^, TMEM173^70^) for CD4+ T cells, then performed gradient-based optimization of the macrophage’s expression profile. Interestingly, the optimization trajectories diverged in the remodeling space, suggesting that rescue objectives drive macrophage modifications along distinct directions (Fig. 4h). To investigate the biological insights of this divergence, we performed limited-move experiments that constrained the optimizer to select only the highest-leverage genes (Fig. 4i, Supplementary Fig. 22,23). The results suggest that the model prioritized a metabolic and adhesive reset for CD4+ T rescue by upregulating FBP1^71^ to reprogram metabolism and downregulating EDIL3^72^ to restore T-cell adhesion. For CD8+ T cell rescue, the model prioritized physical access, identifying the upregulation of collagenases (MMP8^73^) and downregulation of exclusion (LIF^74^) and adhesion (NCAM1^75^) factors.

The final optimized neighborhood modifications (Fig. 4j, Supplementary Fig. 22,23) indicate a comprehensive logic for reactivating adjacent dysfunctional T cells. First, the model prioritized releasing the universal checkpoint brakes specifically by suppressing CD200^76^ alongside ENPP1^77^ and CTHRC1^44^ to simultaneously unlock innate STING signaling and dismantle pathological fibrosis. Second, it triggered a metabolic and innate alarm, requiring the induction of SLC40A1^78^ to restore iron supply and DDX58^79^/NLRP3^80^ to mimic viral sensing. Finally, distinct programs emerged for lineage-specific support: while CD8+ rescue prioritized cytotoxic residency (ITGA1^81^) to anchor effectors, CD4+ optimization paradoxically selected for tissue repair signals (HGF, VSIR^82^). This logic reflects the biological trade-off of helper T-cell rescue, where maximizing stability risked reinforcing a VISTA+ Treg-like niche. Together, these remodeling recipes suggest that activating inflammatory signaling requires the prior dismantling of physical and metabolic barriers. By generating these hypotheses from observational data, TRINUS provides a computational approach for prioritizing candidate interventions for subsequent experimental validation.

In summary, these *in silico* results reveal the distinct interaction syntax underlying the macrophage-mediated immune barrier. By mapping cellular function as a reciprocal negotiation between a cell and its neighbors, TRINUS facilitates the *in silico* engineering of tissue models through the complementary perspectives of forward perturbation and inverse design.

### TRINUS reveals underlying interaction dynamics of developmental systems

To evaluate TRINUS’s capacity for capturing temporal dynamics, we applied it to a mouse embryonic dataset comprising eight spatial transcriptomes from embryonic day E9.5 to E16.5 (Fig. 5a). TRINUS generated prototype embeddings, cell embeddings, and interaction embeddings, decoupling prototype-driven marker expression from interaction-induced deviations (Supplementary Fig. 24-26). For downstream analysis, we clustered the interaction embeddings. While the dominant interactions within a given tissue at a single time point remained relatively stable, they exhibited progressive changes across developmental stages (Fig. 5b, Supplementary Fig. 26).

**Fig 5.**
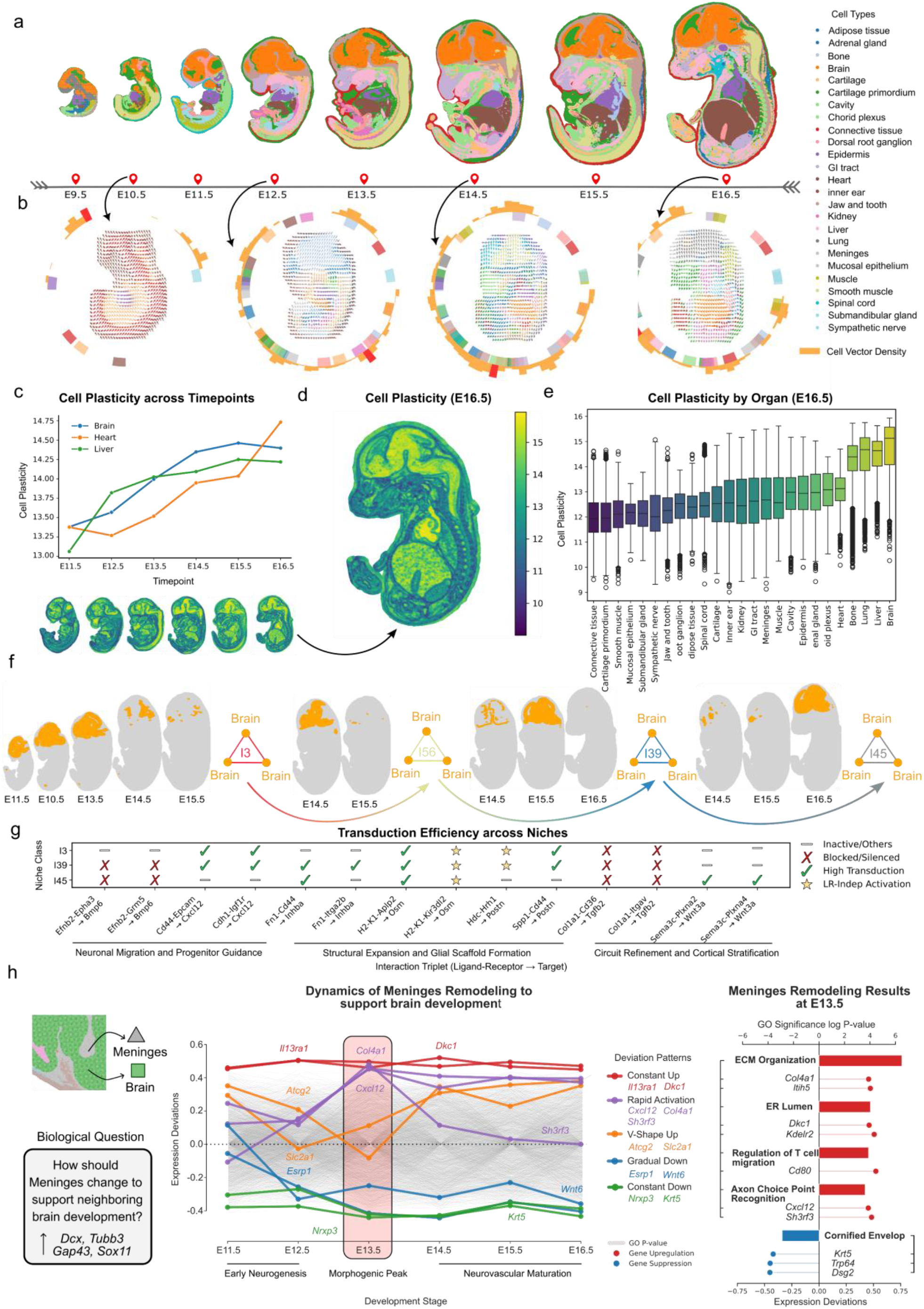
TRINUS reveals interaction dynamics in mouse embryo development. (a) Overview of the mouse embryo dataset across eight developmental time points (E9.5-E16.5). Cell types are color-coded. (b) Interaction compasses for given stages (E10.5, E12.5, E14.5 and E16.5). (c) Cell plasticity mean in brain, heart, and liver from E11.5 to E16.5. (d) Spatial distribution of cell plasticity at E16.5. (e) Cell plasticity mean across organs at E16.5. (f) Dynamics of interaction types during embryonic development. (g) Transduction efficiency analysis across interaction types reveals stage-specific changes in activated ligand-receptor-target (LRT) triplets during embryonic development. (h) In silico niche remodeling of meninges to support the development of brain cells. (Left) Illustration of meninges remodeling (Middle) Trace plot of functional gene contribution during remodeling among different time points. (X-axis) Time points (Y-axis) Expression remodeling in meninges identified. (Right) Detailed gene and GO analysis of most significant genes identified in E13.5 remodeling.

We first quantified cell plasticity using the interaction-induced deviation. Plasticity in brain, heart, and liver increased during embryogenesis, a trend consistent with the progressive integration of cells into increasingly complex functional microenvironments (Fig. 5c). Plasticity also varied substantially across organs. By E16.5, heart, spinal cord, brain, and liver exhibited high plasticity, whereas bone, muscle, and connective tissue were relatively mature and stable, showing lower plasticity (Fig. 5d,e).

Furthermore, we used TRINUS to analyze the dynamics of interaction motifs (Fig. 5f). To interpret functional differences and to demonstrate that our model captures dynamic signaling shifts rather than static expression profiles, we analyzed the transduction efficiency of specific LRT triplets across embryonic brain development (Fig. 5g; I3: E11.5– 13.5; I39: E15.5; I45: E16.5; **Methods**). We observed that interactions driving early neuronal migration, such as Cxcl12 associated signaling axes (involving Cd44 or Cdh1 as annotated targets), exhibited high transduction efficiency during early- and mid-stages, but were correctly identified as inactive by the late stage. Conversely, the model captured late-stage signals. Wnt3a targets, which regulate specific differentiation and late-phase axon guidance^83^, shifted from an inactive state in early development to high transduction exclusively at E16.5 (I45). Furthermore, homeostatic niche maintenance signals, such as Osm pathways^84^, showed sustained activation. The recovery of these distinct temporal profiles indicates that the model distinguishes between transient developmental cues and constitutive environmental signals, effectively inferring the time-dependent correlation between signal potential and target response.

Having shown that TRINUS captures these temporal signaling dynamics, we next asked whether the model could predict the niche remodeling required to support neuronal development across these stages. We performed in silico niche remodeling to determine how the neighboring meningeal niche must change its transcriptional state to actively exploit these windows and facilitate neuronal development by maximizing neuronal migration (Dcx) and differentiation (Tubb3, Gap43, Sox11)^85^ in the neighboring brain (Fig. 5h; **Methods**). Our model revealed that the minimal effort required from the meninges follows a precise, three-stage temporal program (Fig. 5h). During Early Neurogenesis (E11.5-E12.5), the niche prioritized lineage fidelity, characterized by the active suppression of non-permissive epidermal markers (Krt5) and the withdrawal of ectodermal patterning signals (Wnt6^86^). This state transitions into a Morphogenic Pulse peaking at E13.5, where the model predicted a transient, high-magnitude impulse of guidance cues (Cxcl12^87^) and structural anchors (Col4a1^88^) that scaffold neuronal positioning. Subsequently, as the tissue entered Neurovascular Maturation (E14.5-E16.5), the transcriptional prescription pivoted toward metabolic support, driving a robust upregulation of glucose transporters and vascular logic (Actg2,Slc2a1^89^) to fuel the increasing energy demands of the developing cortex.

To further dissect the molecular mechanisms driving the critical E13.5 Morphogenic Pulse, we mapped the prioritized gene gradients to Gene Ontology (GO) terms (Fig. 5h). The analysis reveals a coherent functional architecture: the model identified a requirement for upregulation of ER machinery (Dkc1, Kdelr2) to support the high secretory load required for ‘ECM Organization’ (Col4a1, Itih5). Furthermore, the specific demand for ‘Axon Choice Point Recognition’ (Cxcl12, Sh3rf3) suggests that the model captures not just structural adjacency, but the instructive signaling cues required for proper axonal pathfinding.

Collectively, the alignment of stage-specific interaction motifs or remodeling recipes with the canonical phases of corticogenesis indicates that the learned embeddings could distinguish transient morphogenic signals from constitutive signals. By transforming static transcriptomic snapshots into remodeling trajectories, TRINUS also provides a computational tool to discover the temporal syntax of interactions.

#### TRINUS enhances embeddings from single-cell transcriptomics foundation models

To evaluate the efficacy and adaptability of the NicheLayer module, we integrated it with GeneFormer^90^, a single-cell transcriptomic foundation model. We benchmarked the clustering performance on a kidney VisiumHD dataset, comparing results with and without the NicheLayer (Fig. 6a). The analysis confirms that NicheLayer effectively refines embedding quality. Specifically, precision, recall, and F1 scores for T cells, fibroblasts, and tubular cells improved notably after NicheLayer-based fine-tuning (Fig. 6b,e), demonstrating that adding structural information boosts identification accuracy for specific cell types.

**Fig 6.**
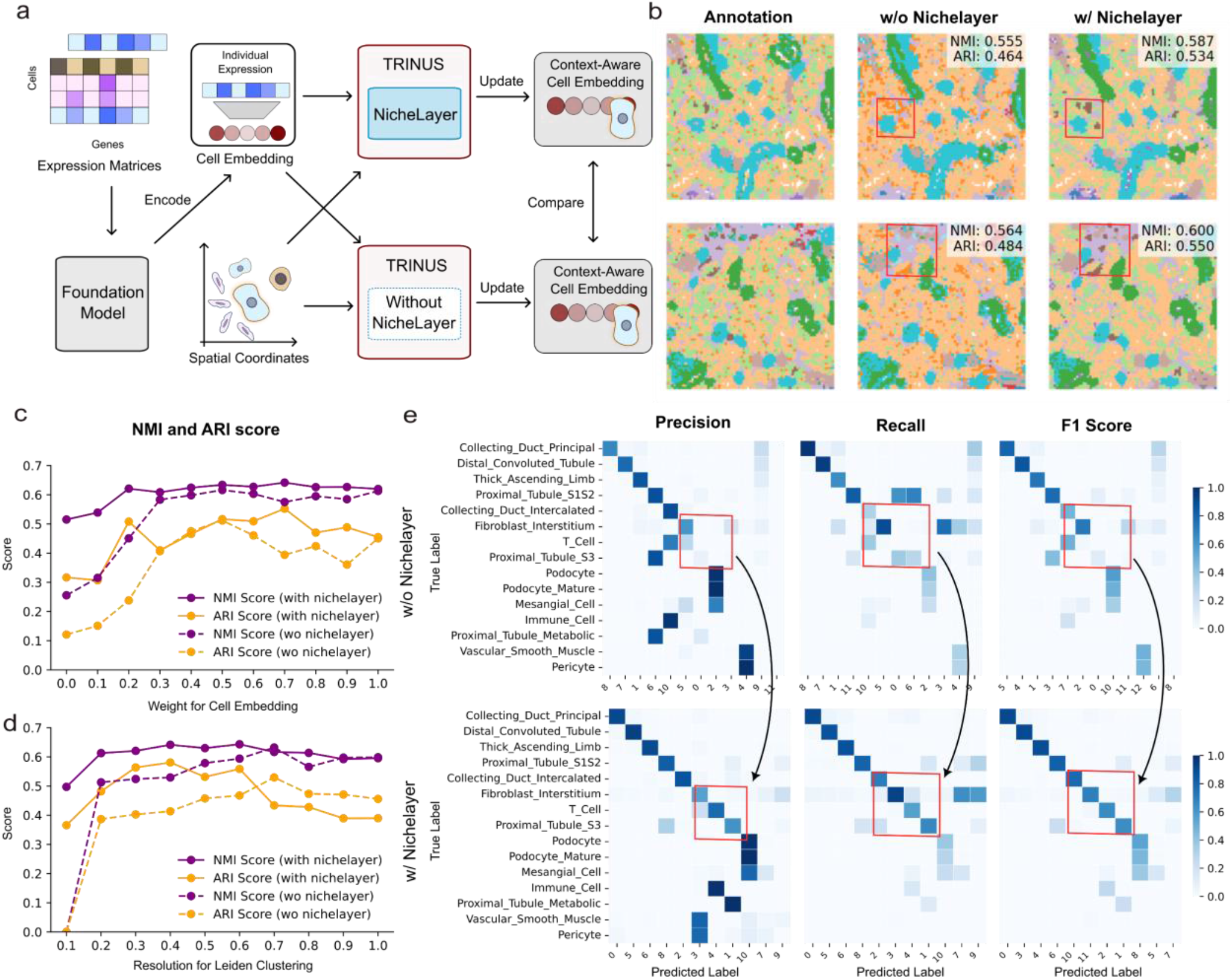
TRINUS enhances cell clustering performance in single-cell foundation models by NicheLayer. (a) Schematic of experimental design: GeneFormer, a spatial transcriptomic foundation model, was evaluated with and without the NicheLayer module on a kidney VisiumHD dataset. (b) Cell clustering performance are improved by NicheLayer, where significantly improved regions are highlighted with red rectangles. (c,d) Evaluation metrics for models with and without NicheLayer under (c) varying weights of cell embedding and interaction embedding, or (d) varying resolutions for Leiden clustering. (e) Precision, recall, and F1 score comparison between models with and without NicheLayer, where significantly improved clusters are highlighted with red rectangles.

Furthermore, we computed similarity matrices with varying weights assigned to cell versus interaction embeddings. The addition of NicheLayer consistently elevated normalized mutual information (NMI) and adjusted rand index (ARI) scores across nearly all weighting schemes. The highest performance was observed at moderate weights (Fig. 6c), and the results are robust around the best resolution for Leiden clustering (Fig. 6d), underscoring that the interaction embeddings contribute valuable information to cell typing alongside the intrinsic cell features.

Finally, we assessed the necessity of external priors by initializing cell embeddings with pre-trained GeneFormer weights (Supplementary Fig. 28). In our experiments, GeneFormer initialization did not lead to improved performance compared to random initialization, and in some cases, metrics showed slight decreases. This observation suggests that TRINUS can learn meaningful cell representations from the data itself, and may not require prior information from other models to achieve its results.

## Discussion

The advent of spatial transcriptomics makes it possible to investigate tissue structure and function. However, it is still difficult to understand how individual cells function through interactions and cooperation. In this work, we introduced TRINUS, a self-supervised model designed to bridge the gap between observational mapping and mechanistic engineering. By allowing the integration between cell states and intercellular interactions, TRINUS demonstrates that tissue function is not merely a sum of cellular parts, but an emergent property of their structural interaction syntax.

Our results suggest that the limitations of current spatial-omics modeling arise from inherent representational bottlenecks in how cellular identity and niche geometry are processed. We have addressed the challenge through two specific resolutions: First, we resolved the identity failure caused by conflated embeddings of smoothing GNNs or the rigid baselines of predictors. By implementing a VQ library and a two-stream information pathway, TRINUS can decouple a cell’s intrinsic lineage (the prototype) from its extrinsic environmental pressure (the deviation) without relying on static population means. Consequently, we could quantify cellular plasticity that reveals how identical lineage states undergo orthogonal reprogramming when placed in distinct hypoxic or exclusion niches. Second, we overcame the geometric failure imposed by the bag-of-cells assumption. Standard message-passing architectures utilize a star topology that treats niche signals as independent, additive weights, blinding these models to the cooperative nature of cell interaction. By introducing triadic multiplicative attention, TRINUS explicitly models the dependencies of cellular signaling networks. This architectural shift enabled the model to perceive the recipe of the niche rather than just the ingredients, leading to the discovery of high-order interaction motifs. Our results show that these motifs are important functional units of tissue organization rather than random cellular aggregates.

In addition, TRINUS enables bidirectional *in silico* experiments. For forward perturbation, we virtually transplant cells to quantify specific biological signals involving resident response (how native tissue reacts to a guest) and transplant adaptation (how the guest adapts to a new niche). For inverse remodeling, we seek the minimal neighborhood modifications required to drive a specific target phenotype change. TRINUS offers three architectural advantages over existing methods in perturbation analysis. Unlike standard GNNs that obscure causality through black-box aggregation, our explicit modeling of interaction residuals ensures that remodeling vectors represent interpretable regulatory shifts. Furthermore, unlike unilateral-based predictors, our model captures reciprocal interactions where targets actively shape their neighbors. Finally, our high-order attention backbone enables global information propagation, avoiding the signal lag and limited receptive fields inherent to message-passing networks, thereby improving state updates during *in silico* experiments.

A primary limitation of TRINUS is the nature of biological supervision: ground-truth labels for single-cell level communications are experimentally inaccessible. Consequently, our training objectives focus on the reconstruction of gene expression, rather than on measuring the physical transmission of molecular signals. Meanwhile, while our *in silico* perturbations align with literature-derived mechanisms, these computational predictions represent hypothesized regulatory links. This hypothesis requires validation through *in vivo* experimental platforms, such as spatial CRISPR screenings or high-throughput functional assays, to confirm that the predicted spatial remodeling yields the desired phenotypic shifts. However, TRINUS could play a complementary role by addressing the challenge of combinatorial complexity within this experimental pipeline. The search space for tissue engineering, the combinatorial possibilities of spatial arrangements and molecular perturbations, is effectively infinite, rendering exhaustive experimental screening impossible. TRINUS functions as a generative hypothesis tool, filtering this infinite space down to a tractable set of high-probability designs. By distinguishing regulatory links from mere spatial correlations, TRINUS provides quantitative possibilities prior to experimental validations.

While the current iteration operates on unimodal spatial transcriptomics, integrating spatial proteomics or epigenomics would allow for a deeper decoupling of factors, separating chromatin-derived intrinsic potential from protein-mediated extrinsic signaling. Furthermore, potential lies in scaling the TRINUS architecture to a spatial interaction foundation model across massive, diverse biological datasets. Such a large-scale model could move beyond local inference to uncover the universal syntax of cellular interactions, distinguishing the conserved, pan-tissue, signaling motifs that define multicellularity from the niche-specific dialects that drive organ-specific functions and pathologies. Overall, TRINUS provides a practical and generalizable tool to investigate the interaction syntax underlying spatial transcriptomics data with the potential of rational tissue engineering design.

## Methods

### TRINUS Overview

To decipher interaction syntax from spatial transcriptomics data, TRINUS represents the tissue as a system of cells whose pairwise interactions are mutually dependent. The model operates on proximal ensemble of cells defined as niches, processing them as complete microenvironments containing both functional cells and cell-cell interactions. For a given niche 𝒩 containing *N* cells, the architecture iteratively refines two distinct representational streams. We introduce Context-Aware Cell Embeddings 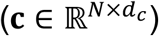 to represent the current state of a cell as influenced by its surroundings and Interaction Embeddings 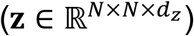 as dense vector representations to encode the molecular and spatial logic between pairs of cells. To resolve the confounding factors of lineage and environment, the model utilizes a Vector Quantized (VQ) bottleneck. This bottleneck maps the context-aware representation onto a library of Context-Free Prototypes 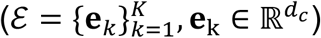 that represents intrinsic states. The network is trained via a composite objective that enforces both accurate individual expression reconstruction and explicit decoupling of observed gene expression into intrinsic prototype markers and interaction-induced deviations. By jointly optimizing these streams within unified and differentiable latent space, TRINUS can also conduct *in silico* simulation, capable of both forward perturbation and inverse niche remodeling.

### Input Data and Embedding Initialization

Let a tissue sample be represented as a set of cells 𝒞, where each cell *i* possesses a gene expression profile **x**_*i*_ ∈ ℝ^*G*^ and spatial coordinates **s**_*i*_ ∈ ℝ^2^. We define a niche 𝒩_*i*_ as the subgraph containing the *N* nearest neighbors centered on cell *i*.

#### Cell Embedding Initialization

The raw expression profile **x**_*i*_ is projected into a latent feature space via a learnable encoder 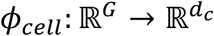. This yields the initial cell representation 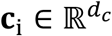. To handle the sparsity and zero-inflation inherent in transcriptomics, we employ a specific encoder architecture dependent on the input modality (standard MLP or foundation-model adapters), with optional masking applied during training to support self-supervised learning.

#### Interaction Embedding Initialization

Representing the social logic of the tissue requires an interaction representation that encodes both molecular potential and spatial proximity. We initialize the interaction embedding 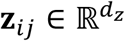 between cells *i* and *j* by concatenating their expression profiles and their spatial relationship:

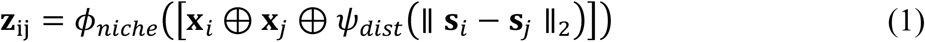

where ⊕ denotes concatenation, *ψ*_*dist*_ is a spatial encoder projecting Euclidean distances into high-dimensional features, and *ϕ*_*niche*_ is a non-linear fusion layer (SwiGLU activated MLP). This initialization ensures that the interaction logic acts as a function of both the sender/receiver molecular states and their physical proximity.

### Niche Layer Architecture

The core computational unit of TRINUS is the NicheLayer, an adaptation of the Evoformer module optimized for spatial transcriptomics. Unlike standard Graph Neural Networks (GNNs) which update node features by aggregating scalar-weighted neighbors (star topology), our architecture maintains and iteratively refines two distinct representational streams: the context-aware cell embedding 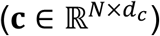 and the interaction embedding 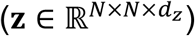. The NicheLayer is designed to enforce geometric and functional consistency within the tissue graph. By stacking these layers, the model enables bidirectional information flow: intrinsic cellular properties shape the interaction logic, while the higher-order structure of the niche dynamically updates cellular identity.

#### High Order Niche Attention

Standard spatial methods treat a cell’s environment as a distinct set of pairwise neighbors (bag of neighbors). This approach fails to capture interaction dependencies, where the functional relationship between cell *i* and cell *j* is heavily dependent on their mutual neighbors. To resolve this, we employ mechanisms^37,38^ that operate explicitly on the cell-cell interactions, enforcing triadic closure as fundamental unit of geometric and social clusters.

To ensure the interaction graph represents a physically and logically consistent signaling space, we apply Triangular Multiplicative Updates. This operation incorporates the inductive bias of transitivity: if cell *i* interacts strongly with *k*, and *k* with *j*, the interaction channel *ij* must be informed by this path. We conduct two symmetric updates (outgoing and incoming) employing element-wise multiplication, which acts as a soft logical gate (i.e., a path exists only if both legs of the triangle are active):

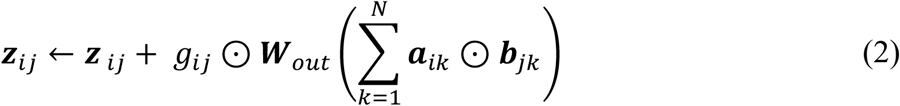

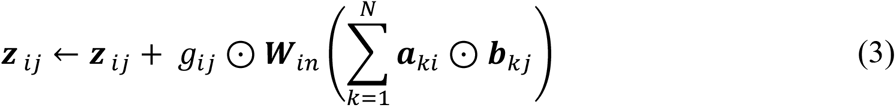

where ⊙ denotes element-wise multiplication, **W** terms are learnable linear projections, and **a, b, g** are gated projections derived from **z**. By summing over the intermediate node *k*, the interaction **z**_*ij*_ accumulates evidence from all possible triangles connecting *i* and *j*, explicitly modeling interaction dependencies and correlations.

While multiplicative updates enforce structural constraints, we employ in addition Axial Attention to refine the *content* of the interaction embedding. This allows an interaction edge *ij* to query the state of other edges sharing a common node. For example, the signaling potential between an immune cell and a tumor cell (*ij*) depends on whether the immune cell is simultaneously engaged with a stromal cell (*ik*). We perform two sequential attention passes over the interaction tensor via starting node attention where edge *ij* attends to all edges *ik* (sharing start node *i*) and ending node attention where edge *ij* attends to all edges *kj* (sharing end node *j*). The update schemes for these scenarios are defined as:

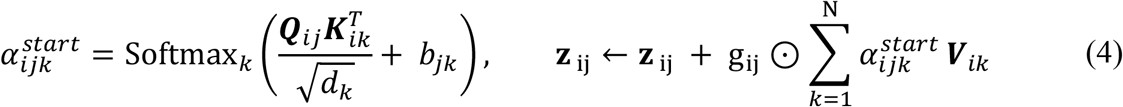

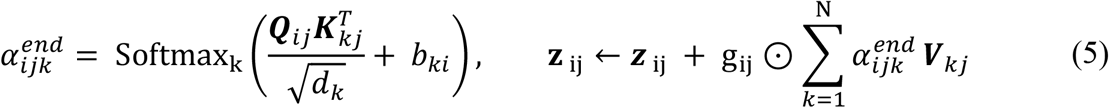

Where ***Q, K***, 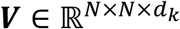 are linear projections of ***z***, and *g*_*ij*_ are learned gate terms, *b*_*ij*_ are learned bias terms. This explicitly enables the model to distinguish distinct topological motifs, such as competitive binding or cooperative signaling clusters, which are invisible to pairwise metrics.

To couple the cellular state with the interaction logic, we employ two specific attention mechanisms that allow the streams to constrain each other.

#### Interaction-to-Identity Flow

The context-aware cell embedding ***c***_*i*_ is updated via interactions in which it participates to reflect its niche. Standard self-attention relies solely on the similarity between cell features (*i* and *j*). We modify this by injecting the learned Interaction Embedding **z**_*ij*_ as a dynamic bias. This ensures the attention mechanism strictly follows the functional topology learned by the interaction stream, rather than Euclidean proximity alone:

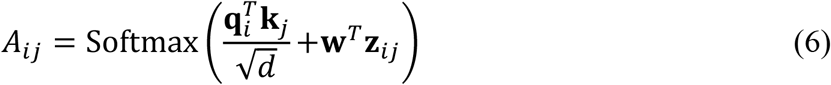

Here, **z**_*ij*_ acts as a learned “molecular distance,” allowing cells to attend strongly to spatially distant but functionally connected partners, while ignoring physically proximal but non-interacting bystanders.

#### Identity-to-Interaction Flow

Conversely, the interaction embedding **z**_*ij*_ is grounded by the instantaneous state of the participating cells. We utilize a Cross-Attention mechanism where the interaction embedding queries the context-aware cell embeddings. This step updates the edge representation **z** based on the plasticity of the nodes, ensuring that as a cell’s state evolves, its interaction potential is updated accordingly.

### Adaptive decoupling via VQ Layer

A fundamental challenge in spatial biology is distinguishing a cell’s intrinsic lineage identity from its plastic measurement driven by environmental candidates. To mathematically decouple these factors without relying on static reference atlases, we utilize a Vector Quantized (VQ) bottleneck^39–41^. We postulate that the observed gene expression **x**_*i*_ is a composite of a discrete intrinsic reference state (Prototype) and a continuous environmental deviation with the Context-Free Prototype ***e***

#### Context-Free Cell Prototypes

The Context-Aware Cell Embedding ***c***_*i*_ (the output of the final Niche Layer) represents the cell’s state *including* spatial influences. To extract the underlying intrinsic identity, we map ***c***_*i*_ to the nearest entry in a learnable codebook 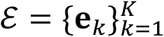, defined as prototypes:

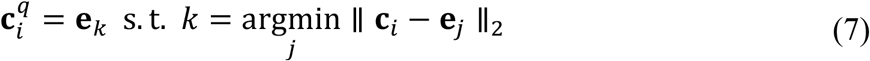

This quantization acts as a rigorous information bottleneck, filtering out unique spatial noise and neighborhood-specific variations, yielding a Context-Free Representation 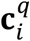. Importantly, this allows the model to define identity dynamically based on the data manifold rather than user-provided labels, avoiding circular dependencies where control cells are already influenced by their niche.

#### Cell Library Updates

To ensure stability during training, the prototype vectors **e**_*k*_ are not updated via gradient descent, but through an exponential moving average^39-41^ of the embeddings ***c***_*i*_ that map to them. Specifically, at each training step, each prototype **e**_*k*_ is updated as a weighted average of its current value and the mean of all embeddings ***c***_*i*_currently assigned to it:

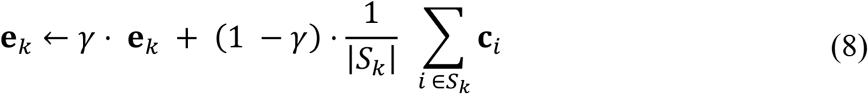

where *S*_*k*_ = { *i* : argmin_*j*_ ∥ ***c***_*i*_ − **e**_*j*_ ∥^2^ = *k* } denotes the set of cells currently assigned to prototype *k*, and *γ* is a momentum coefficient controlling the rate of update. This process ensures that each prototype gradually consolidates toward the consensus transcriptomic state of its assigned population, providing a continuously refined and data-driven reference for each cell lineage.

### Training Objects and Loss Function

The TRINUS model is optimized via a composite objective function designed to ensure that the learned embeddings are biologically meaningful (reconstructive accuracy), spatially coherent (geometric consistency), and robust to technical noise.

#### Hurdle Reconstruction Loss

Spatial transcriptomics data is strictly non-negative and zero-inflated. To deal with these ill-structured data, we model the gene expression using a two-component decoder (*D*^*p*^, *D*^*μ*^) following the insights from Hurdle expression model^91,92^. For any given embedding ***h***, the decoder predicts a tuple (*p* = *D*^*p*^(***h***), *μ* = *D*^*μ*^(***h***)) representing the estimated expression magnitude and the probability of non-zero expression, respectively. We train the probability decoder *D*^*p*^ with binary cross entropy (BCE) loss to predict whether a gene is expressed (**y** = 𝕀_**x**>0_, 𝕀 is the indicator function) and ask the mean decoder *D*^*μ*^ to predict the magnitude of expressed genes. The overall reconstruction loss is defined as:

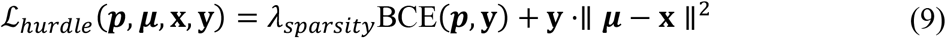

Where *λ*_*sparsity*_ is dataset related coefficient augmenting the importance of zero-inflated counts. This ensures the model learns structural sparsity separately from expression intensity.

#### Decoupling and Reconstruction

We train the network branches using two complementary reconstruction objectives. First, the intrinsic hurdle reconstruction loss ℒ_*cellrecon*_ ensures the context-aware embedding ***c***_*i*_ retains biological fidelity. Second, and crucially, the residual reconstruction forces the interaction embeddings ***z*** to explain the *deviation* of a cell from its standard lineage baseline. We define the interaction effect as the difference between the true observed expression **x**_*i*_ and the decoded Context-Free Prototype 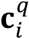 and train the interaction embedding to reconstruct such deviation using the following loss function.

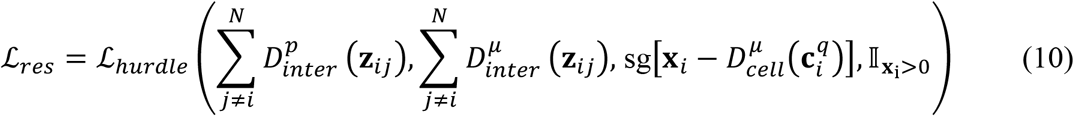

Where sg[] is stop gradient operator that blocks the backpropagation of deviation gradients into the cell embedding branch during the interaction embedding update.

Logically, this treats the predicted intrinsic prototype 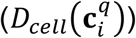 as a step-wise fixed reference anchor. Without this constraint, the model would face a degenerate solution: it could trivially minimize the error by altering the intrinsic prototype to match the total expression, effectively conflating the two signals. By freezing the intrinsic perception during this step, we force the interaction embeddings to explain the information that the intrinsic prototype cannot.

#### Masked Niche Training

To prevent overfitting and enable the model to handle the high dropout rates inherent in spatial transcriptomics, we employ a masked niche training strategy. Unlike standard Masked Language Modeling (MLM) where the objective is to predict the masked token, our approach utilizes masking as a graph-level data augmentation technique. During training, we generate a binary mask vector **m** ∈ {0,1}^*N*^, randomly zeroing out the input features of a subset of cells (simulating technical dropouts or segmentation failures). We also explicitly mask out self-loops in the interaction graph during training to prevent information leakage from a cell to itself via the triangular update paths. The training loss is then computed *only* on the unmasked (visible) cells:

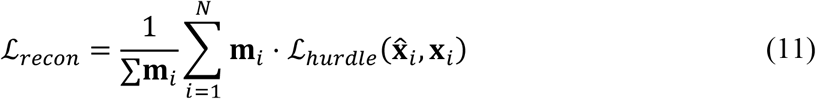

By enforcing reconstruction on the visible nodes despite missing neighbors, the model is compelled to learn robust, distributed interaction patterns rather than relying on brittle, one-to-one correlations.

#### Spatial Constraints

To prevent the interaction embedding from degenerating into a purely semantic vector that ignores physical space, we enforce a spatial bin prediction loss. The continuous Euclidean distances between cells pairwise are discretized into *K* = 4 distinct bins (ranging from immediate contact to distant neighbor). The interaction embedding **z**_*ij*_ predicts this spatial relationship:

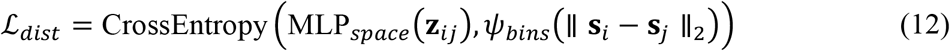

where *ψ*_*bins*_ maps the physical distance to its corresponding class index. This creates a hard constraint ensuring **z**_*ij*_ explicitly encodes spatial proximity.

#### Optional Supervised Fine-Tuning

While TRINUS is fundamentally unsupervised, the architecture supports supervised fine-tuning when cell-type annotations are available. This phase acts as a refinement of the VQ codebook and the interaction logic. We employ an auxiliary focal classification loss (ℒ_*focal*_) with focusing parameter *γ* to handle class imbalance. We apply a classification head to the context-aware embeddings of valid (unmasked) cells to predict their cell type *y*_*i*_:

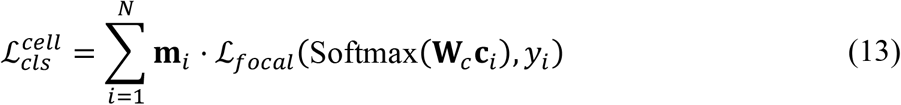

Since the interaction embedding **z**_*ij*_ represents the functional impact of cell *j* upon cell *i*, the aggregate of all outgoing impacts from cell *j* provides a functional fingerprint of *j*’s identity. We predict the cell type of the source node *j* by pooling its influence over the niche:

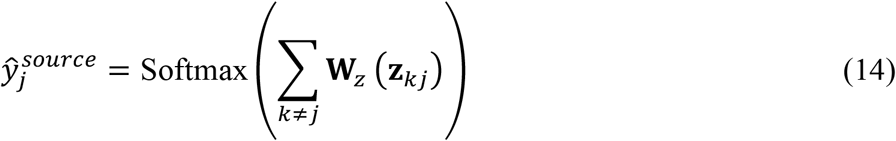

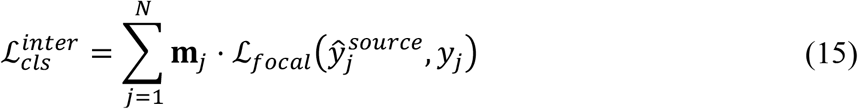

Given these training strategies and loss modules we construct the total loss function as follows:

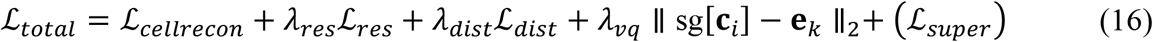

During training we select niche size *N* = 31, *λ*_*res*_ = 10, *λ*_*dist*_ = 0.1, *λ*_*vq*_ = 0.25. And we randomly masked out 30% of genes of each cell and 20% cells within each niche. During training to ensure the smooth training we assign 10 warm-up epochs where *λ*_*res*_ = 0. Optimization is performed using the Adam with an initial learning rate of 10^−4^ and weight decay of 0.01.

### Marker prototype and niche pressure selection

During differential prototype and niche pressure analysis (Fig. 3c), we choose one-versus-rest Wilcoxon rank-sum test from zero-inflation-corrected reconstructed data, following the exclusion of mitochondrial and ribosomal transcripts. To characterize interaction pressures, driver genes were ranked by z-score significance and filtered for a minimum expression fraction (>0.2) in the target niche to ensure robustness, whereas prototype markers were prioritized based on log2 fold change magnitude. The top independent positive and negative drivers per niche were subsequently validated for biological relevance, and a representative subset of these validated markers was selected for visualization via dotplots and heatmaps.

### Adopting Foundation Models

To enhance generalization and leverage prior biological knowledge, TRINUS incorporates adapters for state-of-the-art transcriptomic foundation models, such as GeneFormer^90^ and scFoundation^93^. Instead of training the initial feature encoder from scratch (*f*_*θ*_), we utilize the pre-trained encoders of these models to tokenize processed expression profiles (e.g., ranked values). A linear projection layer maps the varying latent dimensions of these foundation models (*d*_*model*_) to the TRINUS cell embedding dimension (*d*_*cell*_). To accommodate the computational constraints of attention mechanisms while ensuring stable memory usage, input gene sequences are truncated or padded to a fixed token length (e.g., *L* = 256) based on highly variable genes, with strictly masked padding to prevent artifactual attention.

### Interaction Compass: Mapping Vector Fields of Communication

To visualize the spatial logic of tissuewide communication, we developed the interaction compass that would projects high dimensional Interaction Embeddings (**Z**) onto a biologically interpretable 2D vector field as shown in (Fig. 3e, Fig. 5b). Construction requires the continuous embeddings, discretized interaction type labels (derived from clustering), and spatial coordinates (Supplementary Fig. 13).

#### Manifold Projection

We employ Spherical UMAP to project the learned interaction embeddings onto a 1-D cyclic manifold (interval [0,2*π*)). This assigns a continuous interaction azimuth *θ*_*ij*_ to every cell-cell interaction, grouping functionally similar communication patterns into adjacent angular sectors. For each discrete interaction type identified in the tissue, we calculate the circular mean of the azimuths of all edges belonging to that type. This yields a fixed Principal Interaction Direction for each interaction category, ensuring that specific functional dialogues map to consistent angles globally.

#### Vector Field Construction

The tissue slide is tessellated into a global spatial grid. For each grid point, we estimate local densities of each interaction type using Kernel Density Estimation (KDE) over the interaction types.

#### Glyph Visualization

We visualize the dominant communication flows at each grid point using compass glyphs. For each location, we select the top two most abundant interaction types. These are plotted as interaction vectors whose angle corresponds to the *Principal Interaction Direction* of the type (semantic meaning) and magnitude corresponds to the local KDE density (intensity). This partitions the tissue into Interaction Zones defined by the specific combination of active communication channels. Additionally, we create a global Azimuth Histogram to visualize the overall compositional balance of interaction patterns across the sample.

### Cellular Plasticity Evaluation

Given the residual learning and expression decoupling feature of TRINUS, we introduce a quantitative metric for Cellular Plasticity (*PL*). We assume that a cell’s plasticity in a spatial context is reflected by the magnitude of its niche pressure from its intrinsic lineage baseline in response to environmental cues. We then defined the niche pressure as the aggregated output of the interaction decoder for all neighbors *j* acting upon cell *i*. Given the zero-inflated nature of the data, we compute the active deviation magnitude of cell *i* by aggregating the predicted deviation strength ***μ***_*inter,i*_ with the predicted expressed probability ***p***_*inter,i*_.

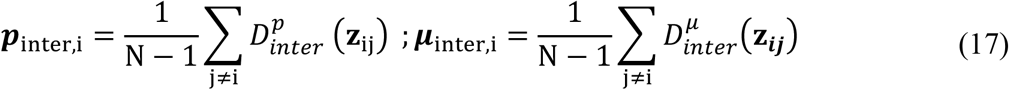

To distinguish functional signaling from background noise, we also apply a hard-thresholding sigmoid gate based on the learned expressed probability. The final plasticity score *PL*_*i*_ is calculated as the L2 norm of the high-confidence vector:

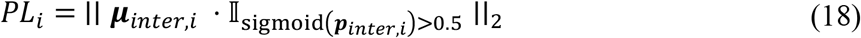

This metric quantifies the adaptation force exerted by the niche on a specific cell. By comparing *PL*_*i*_ across spatial domains, we identify regions where cells are under high environmental pressure versus regions where cells maintain rigid lineage stability (Fig. 3d).

### Communication Efficacy Evaluation

To dissect the context-dependent efficacy of intercellular signaling, we developed a bivariate classification framework integrating structural potential and inferred response (Fig. 3f, Fig. 5g). We prioritized high-confidence signaling axes by retrieving Ligand-Receptor-Target (LRT) triplets from NicheNet^6^, filtering for top-ranked regulatory links supported by literature-curated signaling pathways. For each cell-cell interaction edge, we computed two metrics in gene expression space: communication potential (*CP*): Representing the theoretical capacity for signaling, defined as the product of ligand expression in the sender and receptor expression in the receiver (*CP*_*ij*_ = *L*_*j*_ × *R*_*i*_). Signaling outcome (*O*): Representing the actual cellular response, defined as the model-inferred expression of the target gene 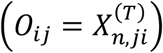. Functional signaling modes were defined through quadrant mapping based on percentile thresholds of the *CP* and *O* distributions. Interactions were categorized into distinct transmission states: High Transduction: High *CP* and high *O*, indicating effective signal propagation.

Silenced/Blocked: High *CP* but low *O*, indicating context-dependent inhibition or receptor desensitization. LR-independent: Low *CP* but high *O*, suggesting alternative activation pathways. Inactive/Others: Low *CP* and low *O* or other states.

### Communication Motifs Discovery

We designed a graph-mining approach to discover and quantify recurrent local interaction patterns from cell spatial neighborhood (KNN) graphs (Fig. 3g, Fig. 5f). Moving beyond isolated edges, we focus on higher-order topological structures formed by multiple edges, specifically analyzing triplets of interaction labels within cellular triangles to detect statistically over-represented cyclic signaling motifs such as A → B → C → A relays. Additionally, we examine star/hub motifs centered on highly connected cells and heterophilic edges between distinct cell types to further characterize potential signaling hubs and dedicated cross-type communication channels. For each motif, we analyze the cell type composition of participating nodes and use occurrence frequency as an initial measure of statistical over-representation, which supports subsequent differential expression analysis to link specific topological configurations to distinct cell states.

### *In Silico* Niche Perturbation

To better understand and probe the syntax of cell-cell interactions, we implemented an *in silico* forward perturbation pipeline (Fig. 4a–g). We probe the collective dynamics of the tissue by computationally determining how the microenvironment responds to, and shapes, specific cellular phenotypes. We define a perturbed condition by virtually transplanting cells of a target type (*C*_*target*_) into the spatial coordinates of a source type (*C*_*source*_), replacing the expression profiles while maintaining physical geometry. To control for the variance introduced by the replacement procedure itself, we conduct a parallel control experiment where *C*_*source*_ locations are filled with different instances of *C*_*source*_ cells. We utilize these conditions for two distinct subproblems: resident response/transplant adaptation analysis.

In resident response analysis we mainly investigate how the surrounding native tissue responds to the new guest. We measure the shift in the interaction induced deviations of the resident neighbors. We define the biological signal Δ_*g,ct*_ as the difference in mean deviations changes between the perturbed (*pert*) and control (*ctl*) groups for each gene *g* and resident cell type *ct*:

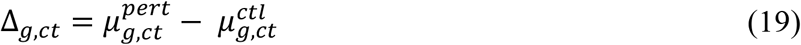

Since the scale of residual is not comparable with full expression profile, we scale such deviation by computing the raw two sample Z-statistics using standard error of the mean (SEM) for both distributions, accounting for the unequal variances of two groups^94,95^.

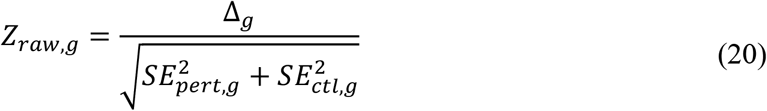

Meanwhile, deep generative models often exhibit over-smoothing, producing output variances lower than biological reality, which might artificially inflate Z-scores. We correct for this by calibrating the test statistic against an Empirical Null derived from the control experiment^96,97^. We estimate the width of the null distribution (*σ*_*null*_)using the robust Interquartile Range (IQR) estimator on the Z-scores of the control residuals^98^. The final adjusted Z-score for gene expression change is defined as

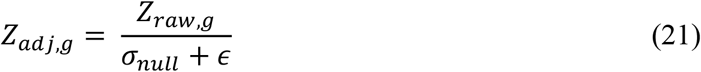

Where *ϵ* is regularization coefficient for numerical stability. This adjustment ensures that reported significance reflects true regulatory shifts (Δ_*g*_) that exceed the background technical and spatial noise of the model.

In transplant adaptation analysis we mainly investigate how transplanted cell adapt to new niches. Here, the perturbation group remains the same (*C*_*target*_ in *C*_*source*_ niche), but the control baseline is defined as investigating *C*_*target*_ cells placed in their native or random *C*_*target*_ niches. This highlights which features of the transplanted cell are plastic responses to the new location versus intrinsic lineage properties. The statistical framework for *Z*_*adj,g*_ computation remains consistent with the resident response analysis.

### *In Silico* Niche Remodeling

The explicit modeling of interactions via dense embeddings enables TRINUS to perform *in silico* niche remodeling: calculating the minimal neighborhood expression modifications required to drive a specific phenotypic transition (Fig. 4h–j, Fig. 5h). Given a central target cell with expression *x*_*target*_ and a desired set of marker genes to modulate, we freeze the model weights and optimize the input gene expression vectors of the neighboring source cells (*x*_neigh_). During remodeling we minimize such optimization objective to drive the target cell toward the desired state, subject to biological constraints:

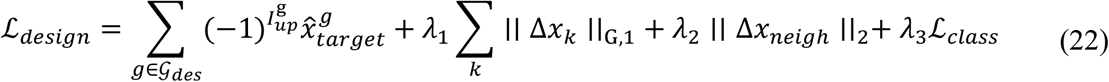

Such loss function describes (1) Directional Objective: The first term maximizes (*I*_*up*_ = 1) or minimizes (*I*_*up*_ = 0) the interaction induced reconstructions of gene *g* in desired gene set 𝒢_*des*_. (2) Sparsity Constraints (*λ*_1_)^36^: A Group-L1 penalty enforces sparsity on the modified genes, ensuring the designed intervention requires changing as few genes as possible. (3) Magnitude (*λ*_2_): An L2 penalty limits the total volume of expression change to biologically plausible ranges. (4) Lineage Constraint (*λ*_3_): A classification loss ensures the optimized neighbors remain within the manifold of their original cell type identity, preventing the generation of unrealistic Frankenstein cells.

### Benchmark for context-aware cell embedding

For context-aware cell embedding, we used four standard evaluation metrics to quantify the consistency between the annotated ground truth and the model’s predictions. Assume *Y* = {*y*_1_, *y*_2_, …, *y*_*n*_} denotes as the ground truth (i.e. cell type label), and 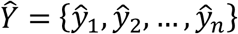 denotes as the model’s prediction. Details of implementing or applying benchmarking methods can be found in **Supplementary Notes**.

Normalized Mutual Information (NMI). NMI quantifies the mutual information between *Y* and 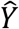, normalized by their entropies:

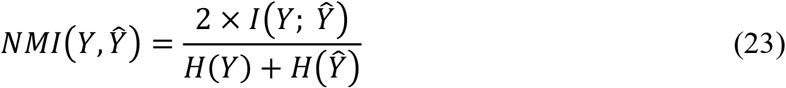

where 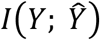 denotes as the mutual information between *Y* and 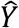:

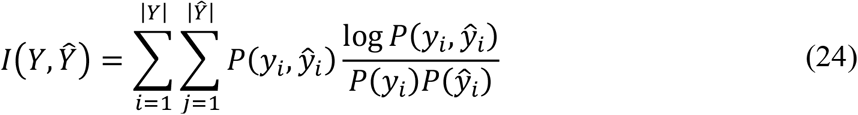

and *H*(*Y*) and 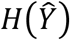 denote as the entropy of *Y* and 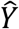 respectively:

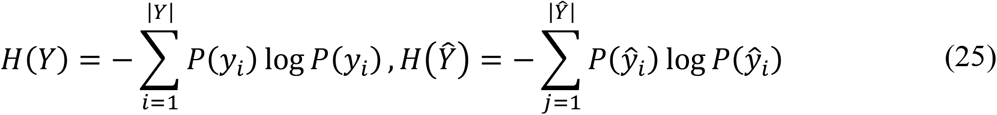

Adjusted Rand Index (ARI). ARI measures the similarity between *Y* and 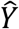, correcting for chance agreement:

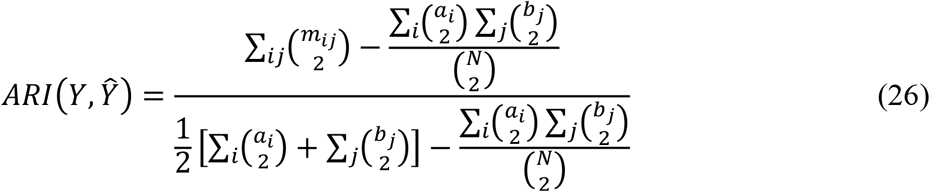

where *M* = [*m*_*ij*_] denotes as the confusion matrix. *a*_*i*_ = ∑_*j*_ *m*_*ij*_, *b*_*j*_ = ∑_*i*_ *m*_*ij*_, *N* = *a*_*i*_ = ∑_*ij*_ *m*_*ij*_.

Homogeneity score. Homogeneity evaluates whether each predicted cluster contains only members of a cell type:

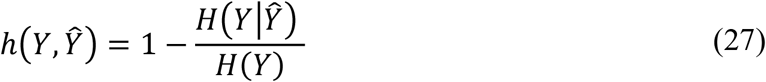

where *H*(*Y*) denotes as the entropy of *Y*, and 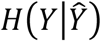 denotes as the conditional entropy of 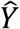 given *Y*:

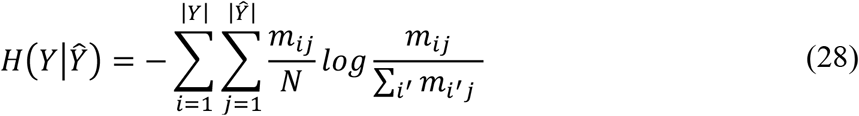

Completeness score. Completeness assesses whether all members of a given cell type are assigned to the same predicted cluster:

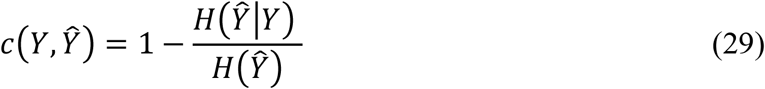

where 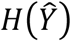 denotes as the entropy of 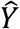, and 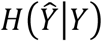 denotes as the conditional entropy of *Y* given 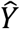:

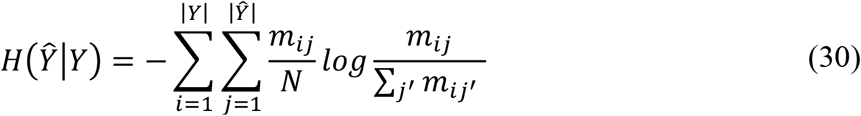

All metrics are positively correlated with the quality of the model’s predictions, and were computed using scikit-learn. Before benchmarking, cells with missing type labels were excluded to ensure evaluation accuracy. For each method, we calculated the above metrics between the ground truth labels and the clustering derived from the cell embeddings (where applicable).

### Benchmark for interaction embedding

Interaction embeddings encode cell–cell communication signals, which are conceptually distinct from both cell-type identity and spatial region. To assess whether clustering based on these embeddings reflects functionally relevant organization, we selected four key biological processes and compiled a gene set ℳ_*g*_ = {*g*_1_, *g*_2_, …, *g*_*K*_} for each process. Given a clustering result with M clusters (*C*_1_, *C*_2_, …, *C*_*m*_) and gene expression *X* = [*x*_*ik*_], we defined a biological consistency score (BCS) as follows:

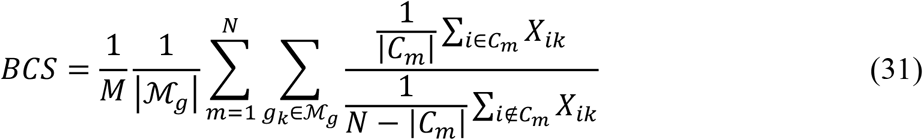

The value *BCS* > 1 indicates that the clustering significantly enriches genes from the predefined functional set M*M*, implying that the derived clusters align with known biological processes and cell-type functions. Other details of implementing or applying benchmarking methods can be found in **Supplementary Notes**.

### Spatial-colocalization calculation

To quantify the spatial correlation between distinct epithelial cell states, we computed a co-localization matrix based on Bivariate Moran’s I. First, we constructed a spatial k-NN adjacency matrix 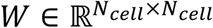, where *w*_*ij*_ ∈ {0, 1} indicates whether cells i and j are neighbors. Then we defined a binary indicator matrix 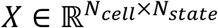, where *x*_*ip*_ ∈ {0, 1} denotes whether cell i belongs to state p. The Bivariate Moran’s I between cell states i and j was then calculated as follows:

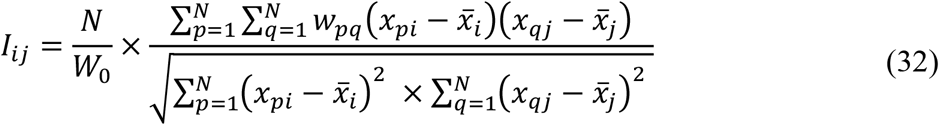

where N denotes as the total cell counts. *I*_*ij*_ > 0 indicates positive spatial co-localization between cell states i and j, whereas *I*_*ij*_ < 0 suggests spatial segregation or negative association.

## Data Availability

Dataset investigated in this work are all publicly available. VisiumHD CRC dataset is available at https://www.10xgenomics.com/cn/datasets/visium-hd-cytassist-gene-expression-libraries-of-human-crc-v4, annotation for this dataset is available at paper S2Omics^99^. Xenium 5k OV and COAD dataset is available at SPATCH ^100^(http://spatch.pku-genomics.org/#/dataset/xenium). Stereo-seq Embryogenesis dataset is available at https://db.cngb.org/search/project/CNP0001543. VisiumHD Kidney dataset is available at https://www.10xgenomics.com/cn/datasets/visium-hd-cytassist-gene-expression-libraries-human-kidney-ffpe.

## Code Availability

The code supporting the findings of this study will be made publicly available upon acceptance of the manuscript following peer review.

## Acknowledgments

This work was supported by the National Natural Science Foundation of China (No. T2321001 to L.Z. and P.Z., 12225102 to L.Z., and 12288101 to L.Z. and P.Z.) and the National Key Research and Development Program of China 2024YFA0919500 (to L.Z.).

## Author Contributions

L.Z., Q.N., P.Z. and Q.G conceived the project. Q.G., W.Z. and Z.Z. designed the model and conducted all analysis. Q.G., W.Z., Z.Z., L.Z., P.Z. and Q.N. addressed the draft. L.Z., P.Z. and Q.N. supervised the research.

